# Uncovering biomarkers during therapeutic neuromodulation with PARRM: Period-based Artifact Reconstruction and Removal Method

**DOI:** 10.1101/2020.10.02.322743

**Authors:** Evan M. Dastin-van Rijn, Nicole R. Provenza, Jonathan S. Calvert, Ro’ee Gilron, Anusha B. Allawala, Radu Darie, Sohail Syed, Evan Matteson, Gregory S. Vogt, Michelle Avendano-Ortega, Ana C. Vasquez, Nithya Ramakrishnan, Denise N. Oswalt, Kelly R. Bijanki, Robert Wilt, Philip A. Starr, Sameer A. Sheth, Wayne K. Goodman, Matthew T. Harrison, David A. Borton

## Abstract

Advances in device development have enabled concurrent stimulation and recording at adjacent locations in the central nervous system. However, stimulation artifacts obscure the sensed underlying neural activity. Here, we developed a novel method, termed Period-based Artifact Reconstruction and Removal Method (PARRM), to remove stimulation artifacts from neural recordings by leveraging the exact period of stimulation to construct and subtract a high-fidelity template of the artifact. Benchtop saline experiments, computational simulations, five unique *in vivo* paradigms across animal and human studies, and an obscured movement biomarker were used for validation. Performance was found to exceed that of state-of-the-art filters in recovering complex signals without introducing contamination. PARRM has several advantages: it is 1) superior in signal recovery; 2) easily adaptable to several neurostimulation paradigms; and 3) low-complexity for future on-device implementation. Real-time artifact removal via PARRM will enable unbiased exploration and detection of neural biomarkers to enhance efficacy of closed-loop therapies.

**Summary:** Online, real-time artifact removal via PARRM will enable unbiased exploration of neural biomarkers previously obscured by stimulation artifact.

## Introduction

The development of closed-loop electrical neuromodulation therapies, for example adaptive Deep Brain Stimulation (aDBS) and adaptive Spinal Cord Stimulation (aSCS) would revolutionize the efficacy of neurostimulation therapies for treatment of many disorders, including Parkinson’s Disease (PD) (*1–3*), Epilepsy (*4, 5*), Essential Tremor (*6*), Obsessive Compulsive Disorder (OCD) (*7*), Treatment Resistant Depression (TRD) (*7, 8*), and chronic pain (*9*). While significant advances have been made in biomarker identification in PD and Epilepsy (*5, 10*), there is no definitive biomarker for a single mental disorder, including OCD and TRD, or chronic pain (*7–9*). Biomarker identification and development of an adaptive neurostimulation system requires a hardware platform that is capable of simultaneous sensing and stimulation. This is particularly challenging when the neural signal of interest originates in or nearby the stimulation target as the amplitude of stimulation therapy is typically several orders of magnitude greater than the amplitude of signals of interest in the brain and spinal cord. Therefore, recordings for adaptive control are heavily contaminated by high amplitude, high frequency stimulation artifact (*11*). In order to extract the underlying neural signatures of disease state, it is necessary to remove the stimulation artifact.

Typically, high frequency artifacts are removed using a lowpass filter, however, limited sampling rates of existing implantable DBS and SCS devices and aliasing of stimulation pulses into low frequencies render lowpass filters ineffective. Existing stimulation artifact removal methods robust to aliasing typically fall into one of three categories: signal reconstruction via deletion and interpolation, decomposing and subtracting components of the signal related to the artifact, and subtracting a template of the artifact at each stimulation pulse. Deletion and interpolation is not effective when artifact duration is long, and is not ideal due to signal loss over the duration of each artifact (*11*). Signal decomposition methods have shown some success, but require a large set of recording channels to be effective (*12*). Template subtraction methods have proven to be successful, however they rely on accurate detection of each stimulation pulse (*13, 14*). Existing methods for identifying individual stimulation pulses in recorded data (e.g. thresholding) are not robust to low sampling rates, the presence of other spurious high amplitude artifacts, or stimulation artifacts with broad peaks(*15*). To our knowledge, there are currently no methods effective at removing stimulation artifacts from LFP recordings sampled at less than twice the frequency of stimulation without contaminating the underlying neural signal, thus greatly hindering control signal identification.

To overcome the challenges in removing periodic stimulation from neural recordings, we have developed a novel artifact removal method, Period-based Artifact Reconstruction and Removal Method (PARRM), to remove high frequency stimulation artifact in low and high-resolution LFP recordings. We demonstrate that PARRM has superior performance to existing, state-of-the-art filters in saline experiments, computer simulations, and five unique *in vivo* recording paradigms. Finally, we demonstrate that PARMM enabled the recovery of a previously obscured biomarker in Parkinson’s Disease participants and could be implemented online to perform real-time biomarker detection (Figure 1).

**Figure 1:**
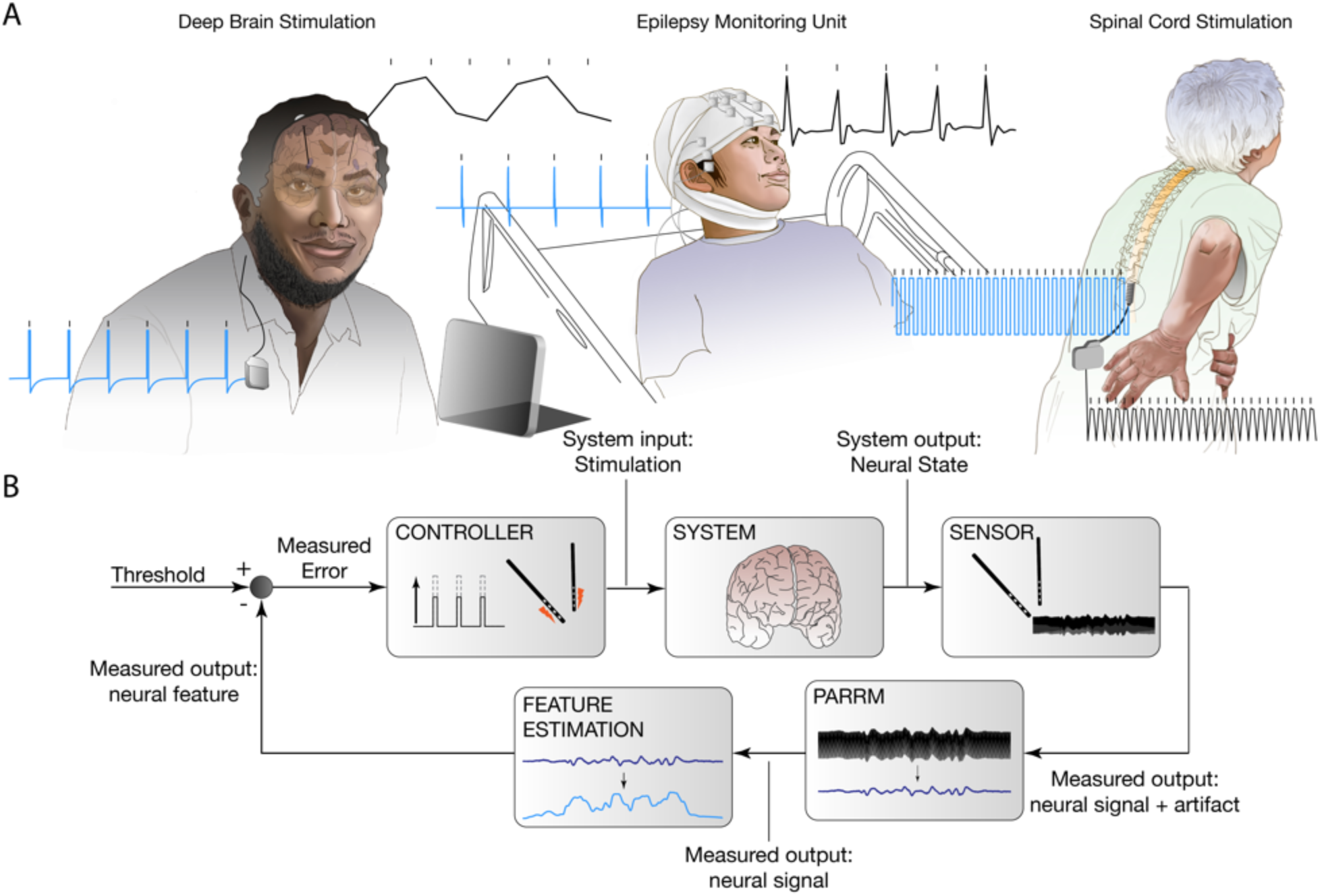
PARRM will enable biomarker detection during ongoing neurostimulation to enhance efficacy of closed-loop neuromodulation. (a) DBS applied at 150 Hz via the Activa PC+S for treatment of refractory OCD (top), DBS applied at 120 Hz in an Epilepsy Monitoring Unit-like (EMU-like) scenario for treatment of TRD, and SCS applied at 50 Hz for treatment of chronic pain. Blue trace shows theoretical injected DBS waveform and black trace shows DBS waveform sampled in vivo at 200 Hz, 2 kHz, and 30 kHz, via the Activa PC+S, Blackrock Cerebus, and Ripple Nomad, respectively. (b) Control policy for closed-loop DBS. Electrodes in the brain sense neural signal and artifact. Real-time artifact removal via PARRM removes neural artifact without contaminating the underlying neural signal, enabling feature estimation for the closed-loop control of stimulation amplitude to relieve symptoms.

## Results

### Design of PARRM

PARRM subtracts an estimate of the stimulation artifact at each time bin from the recorded signal at that time bin. The artifact estimate is formed by averaging the recorded signal at other time bins that are close to the current time bin in *both* time and stimulation phase. The artifact is presumed to be roughly identical for all of these time bins. Averaging reduces the influence of brain signals and additional sources of noise, so that the estimate is primarily artifact. This process can be implemented as a linear filter (i.e., a weighted average using a sliding window). PARRM needs a precise estimate of the stimulation period relative to the sampling rate. Slight inaccuracies in device system clocks can necessitate using a data-driven method to determine this period, which is done by finding the period that, when the data are divided into epochs the length of one period and overlapped, the samples will consolidate around the shape of the high-resolution artifact waveform. The complete process of data-driven period finding, artifact estimation, and signal reconstruction is illustrated in Figure 2.

**Figure 2:**
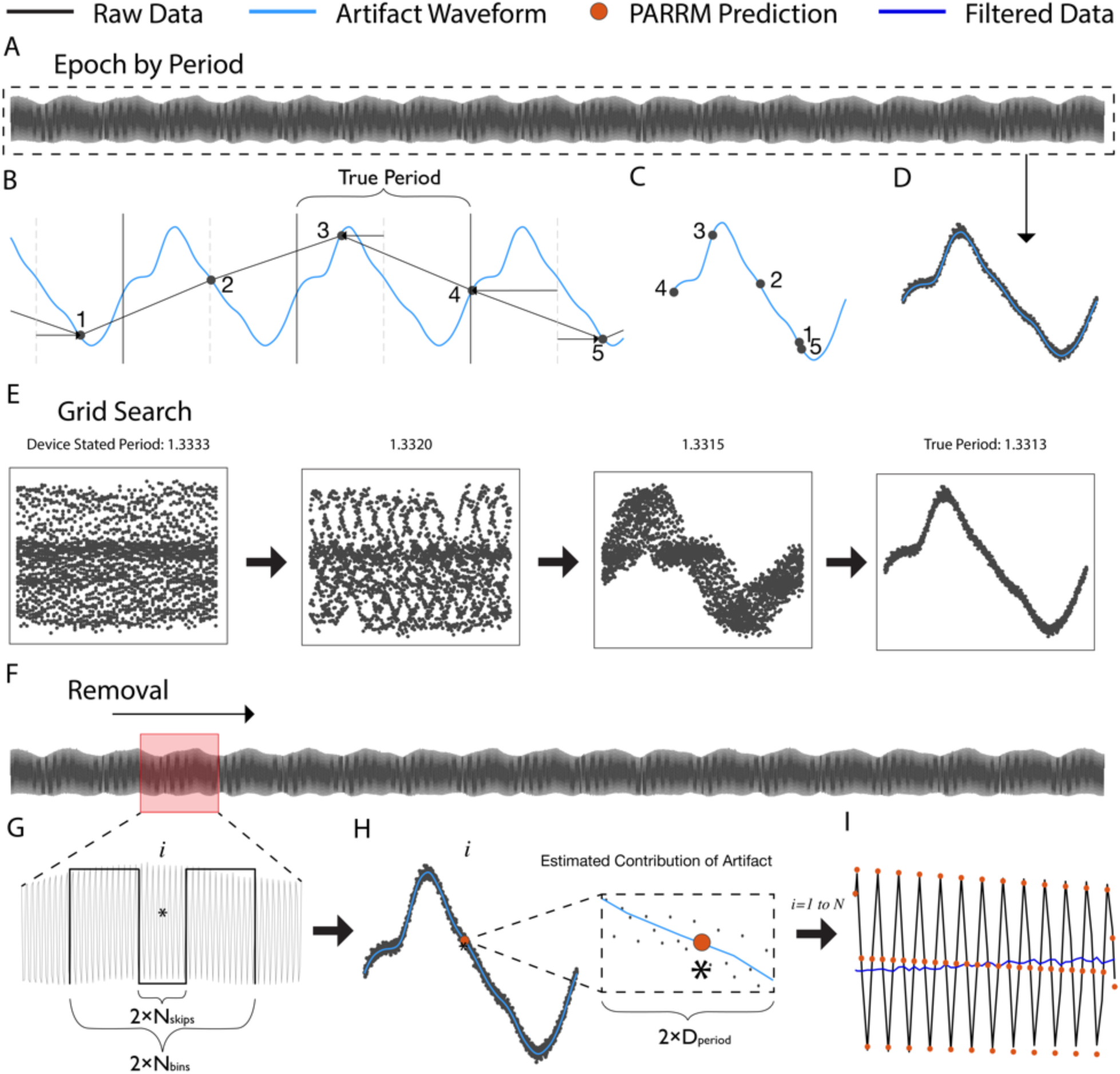
Illustration of stimulation period determination, template reconstruction, and template subtraction via PARRM. (a) Entire LFP recording sampled at 200 Hz (black) is used to identify the true period. (b) An illustration of a 5-sample snippet of the LFP recording divided into epochs using the true period and overlaid with the high-resolution waveform (light blue). Black points indicate individual raw LFP samples. (c) The epochs for all 5 samples are overlaid on the timescale of the true period. (d) When all epochs in the recording are overlaid using this procedure, all samples consolidate around the shape of the high-resolution artifact waveform on the timescale of the true period. (e) The period suggested by the device sampling and stimulation rates is inexact and does not result in a consolidated waveform. Using a grid search centered around the stated period, a series of periods are evaluated to find the true period that produces the most consolidated samples. (f) A sliding window is applied to the entire recording to estimate the contribution of the stimulation artifact at each sample. (g) For each window, a rectangular kernel (length*N*_bins_), ignoring the center (length *N*_skips_) is used to estimate the value of the artifact at each sample of interest *i* (asterisk). (h) Samples within a distance, *D*_period_, on the timescale of the artifact period are averaged together to produce the estimate of the amplitude of the artifact (orange point) at sample *i*. (I) The estimated value of the artifact is then subtracted at each sample over the entire recording to recover the signal of interest (dark blue).

### PARRM recovers simple sinusoidal signals in saline

PARRM was used to remove the DBS artifact and recover the underlying injected signal and noise in saline. In artifact free (DBS off) recordings, both the 10 and 50 Hz injected signals are clearly visible both in the frequency and time domains prior to signal offset (Figure 3a, 3b). When stimulation is turned on, high amplitude artifacts are visible in the frequency domain at 0 and 50 Hz, obscuring the 50 Hz injected signal but not the 10 Hz signal. In the time domain, both the 10 Hz and 50 Hz signals are obscured (Figure 3c, 3d). Following filtering using PARRM, the effects of stimulation are removed in both the frequency and time domains (Figure 3E, 3F). In the case of the 50 Hz signal, this is achieved despite the artifact being aliased to the same frequency as the injected signal.

**Figure 3:**
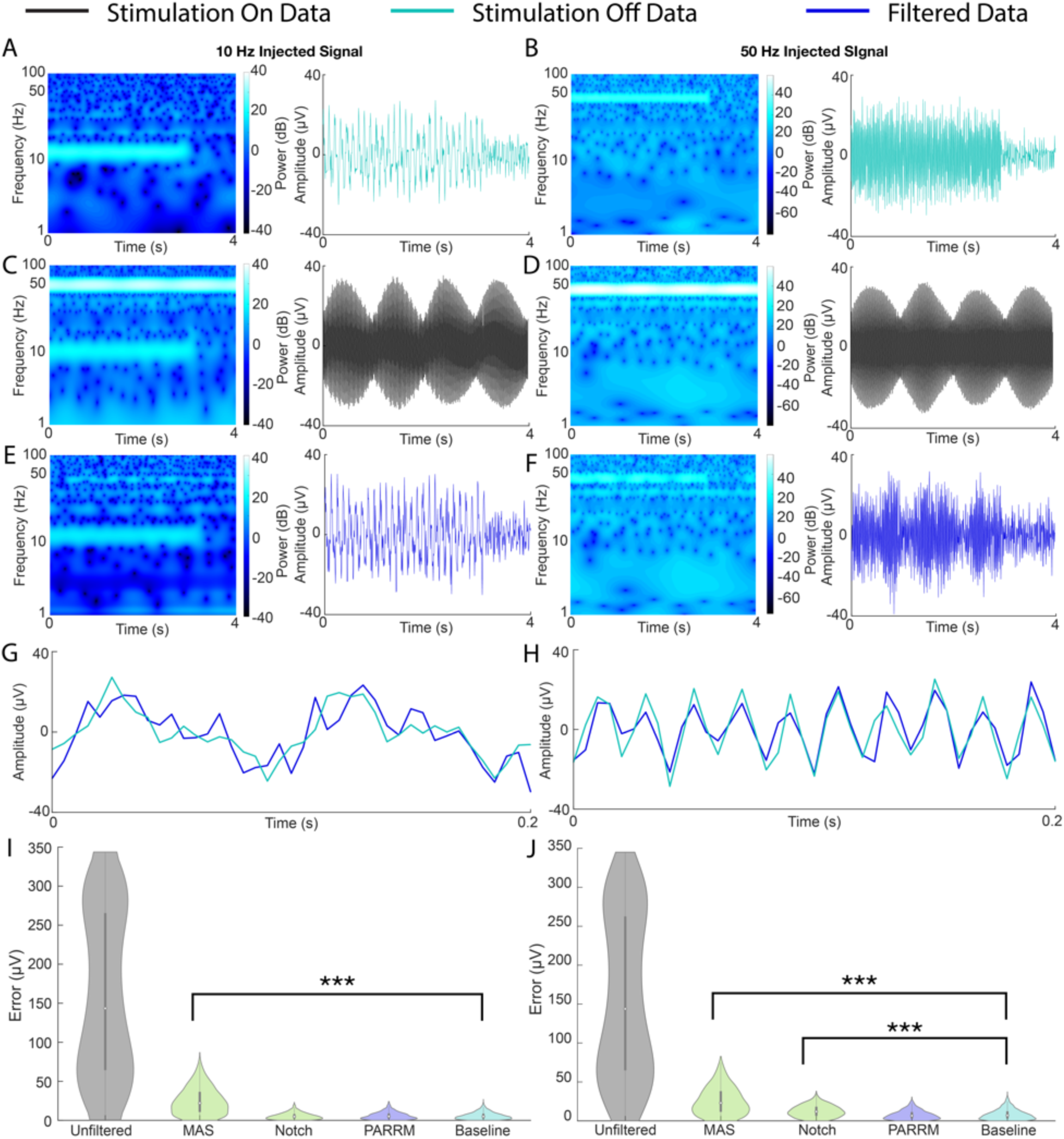
PARRM effectively recovers sinusoidal signals at frequencies separate from and coincident with the aliased artifact. (a, b) Spectrogram and time-voltage series of 10 Hz and 50 Hz sinusoidal signals injected into saline sampled at 200 Hz with stimulation off. (c, d) Spectrogram and time-voltage series of 10 Hz and 50 Hz sinusoidal signals injected into saline sampled at 200 Hz during concurrent 150 Hz stimulation. (e, f) PARRM filtered spectrogram and time-voltage series of 10 Hz and 50 Hz sinusoidal signals injected into saline sampled at 200 Hz during concurrent 150 Hz stimulation. (g, h) A 0.2 second snippet of PARRM filtered and artifact free time-voltage series of 10 Hz and 50 Hz sinusoidal signals injected into saline sampled at 200 Hz during concurrent 150 Hz stimulation. (i, j) Evaluation of filter performance based on time domain absolute error between artifact free and filtered 10 Hz and 50 Hz injected signals sampled at 200 Hz during concurrent 150 Hz stimulation. Asterisks indicate significant differences from absolute errors on the order of baseline noise (Wilcoxon ranksum, ***: *p<0.0005*).

The similarity between the artifact free and filtered signals is visually apparent in 0.2 seconds of data for both the 10 and 50 Hz injected signals (Figures 3g, 3h). We then quantified filter performance by comparing the distribution of absolute errors between artifact free signals and unfiltered, MAS filtered, notch filtered, and PARRM filtered signals to a baseline noise recording (no stimulation, no injected signal) (Figure 3i, 3j). Filtering using MAS did not reduce the error to the level of baseline for either injected signal (p<0.0005). While effective at 10 Hz (p>0.05), Notch filtering removed the injected signal along with the artifact leading to a large reduction in error yet still significantly different from baseline (p<0.0005). For both the 10 and 50 Hz injected signals, PARRM outperformed the other methods with no significant difference (p>0.05) from baseline, indicating that the remaining errors were expected due to noise in saline.

### PARRM recovers complex, multi-frequency signals in computer simulations

Having shown that PARRM is effective for recovering simple sinusoidal signals recorded in saline, next we sought to compare the method’s performance to a series of state-of-the art filters in recovering more complex, injected, chirp signals for simulated data (S. Fig. 4). When all chirps were averaged, PARRM recovered a signal with minimal distortion and noise in the time domain at both low and high sampling rates, unlike MAS and matched filters (Figure 4a left, 4b left). Additionally, PARRM showed no significant differences in the frequency domain at either sampling rate (Figure 4a right, 4b right). This was true even at frequencies affected by artifact where other filters either over (notch) or under (MAS and matched) filtered. Lastly, PARRM had a relative root mean squared error (RRMSE) close to one for both sampling rates indicating effective signal recovery on a single trial basis exceeding performance of the Hampel filter (Figure 4c, 4d). For all three metrics, PARRM exceeded performance of all other filters for both low and high sampling rates.

**Figure 4:**
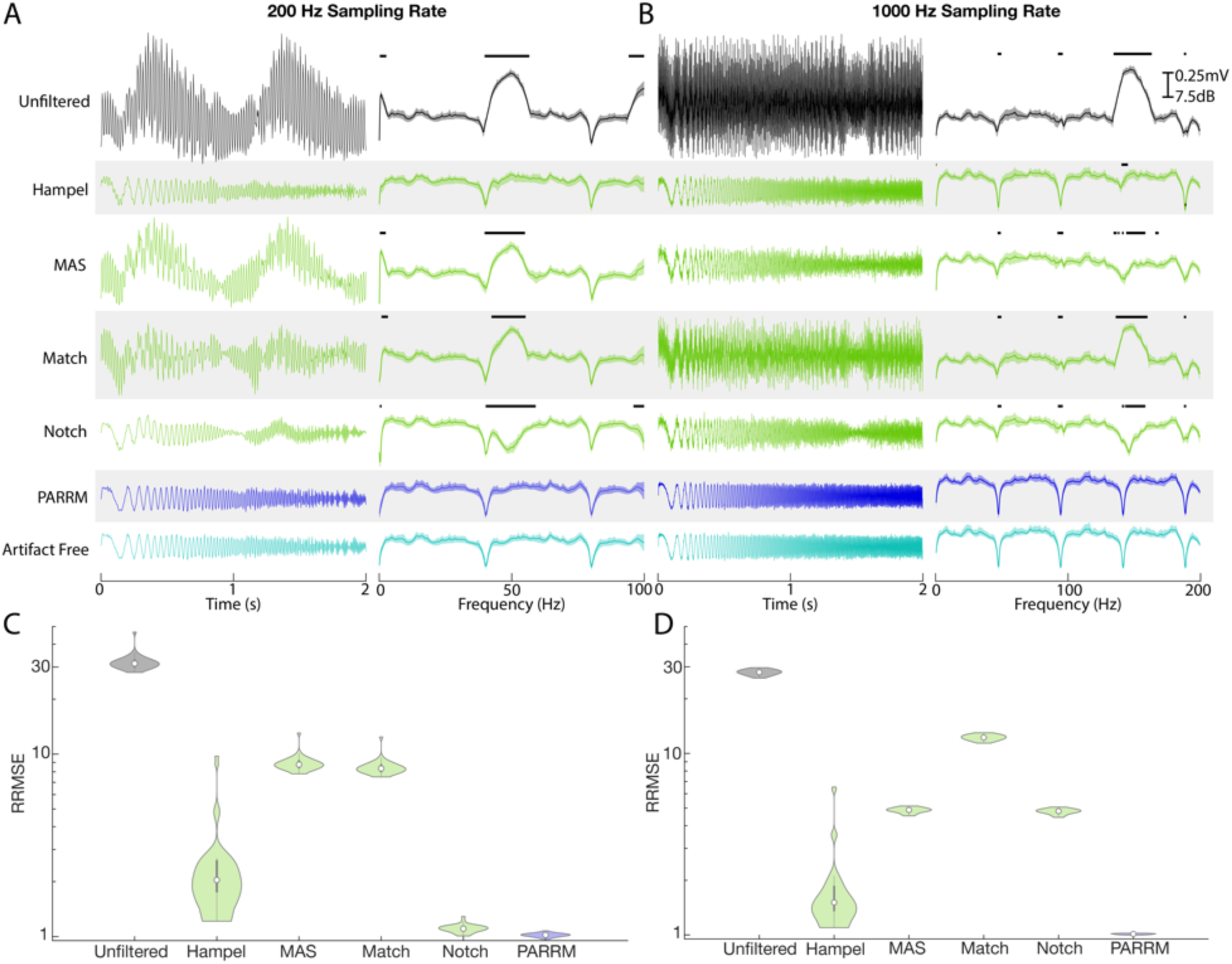
PARRM performance exceeds state of the art filters for nonstationary signals at low and high sampling rates in simulated data. (a) Averaged time-voltage series and windowed power spectral density of 30 simulated linear chirps (0 Hz to 100 Hz, 2 second duration, variable separation) during concurrent 150 Hz stimulation for unfiltered, Hampel filtered, MAS filtered, Match filtered, Notch filtered, PARRM filtered, and artifact free recordings sampled at 200 Hz. Black solid bars indicate significant difference from artifact free signal (2-Sample t-test, *p < 0.05*). (b) Average time-voltage series and average windowed power spectral density of 30 simulated linear chirps (0 Hz to 200 Hz, 2 second duration, variable separation) during concurrent 150 Hz stimulation for unfiltered, Hampel filtered, MAS filtered, Match filtered, Notch filtered, PARRM filtered, and artifact free recordings sampled at 1000 Hz. Black solid bars indicate significant difference from artifact free signal (2-Sample t-test, *p < 0.05*). (c, d) Evaluation of filter performance based on time domain relative root mean squared error (RRMSE: ratio between MSE of artifact free vs. theoretical chirp to MSE of filtered vs. theoretical chirp) of simulated chirps during concurrent 150 Hz stimulation sampled at 200 Hz and 1000 Hz.

Next, a parameter sweep was performed to test the effect of varying chirp length (1-10 s), amplitude (0.5-5 V), pulse width (30-180 µV), and frequency (80-180 Hz) on PARRM performance, measured by RRMSE and Relative R Ratio (S. Fig. 5). Effects for chirp length and pulse width were all within the margin of error. RRMSE and Relative R Ratio increased for increasing stimulation amplitude. RRMSE and Relative R Ratio decreased for stimulation frequencies above 100 Hz. All changes were at most 8% different from baseline indicating that PARRM performed well for a wide range of stimulation parameters and recorded signals.

**Figure 5:**
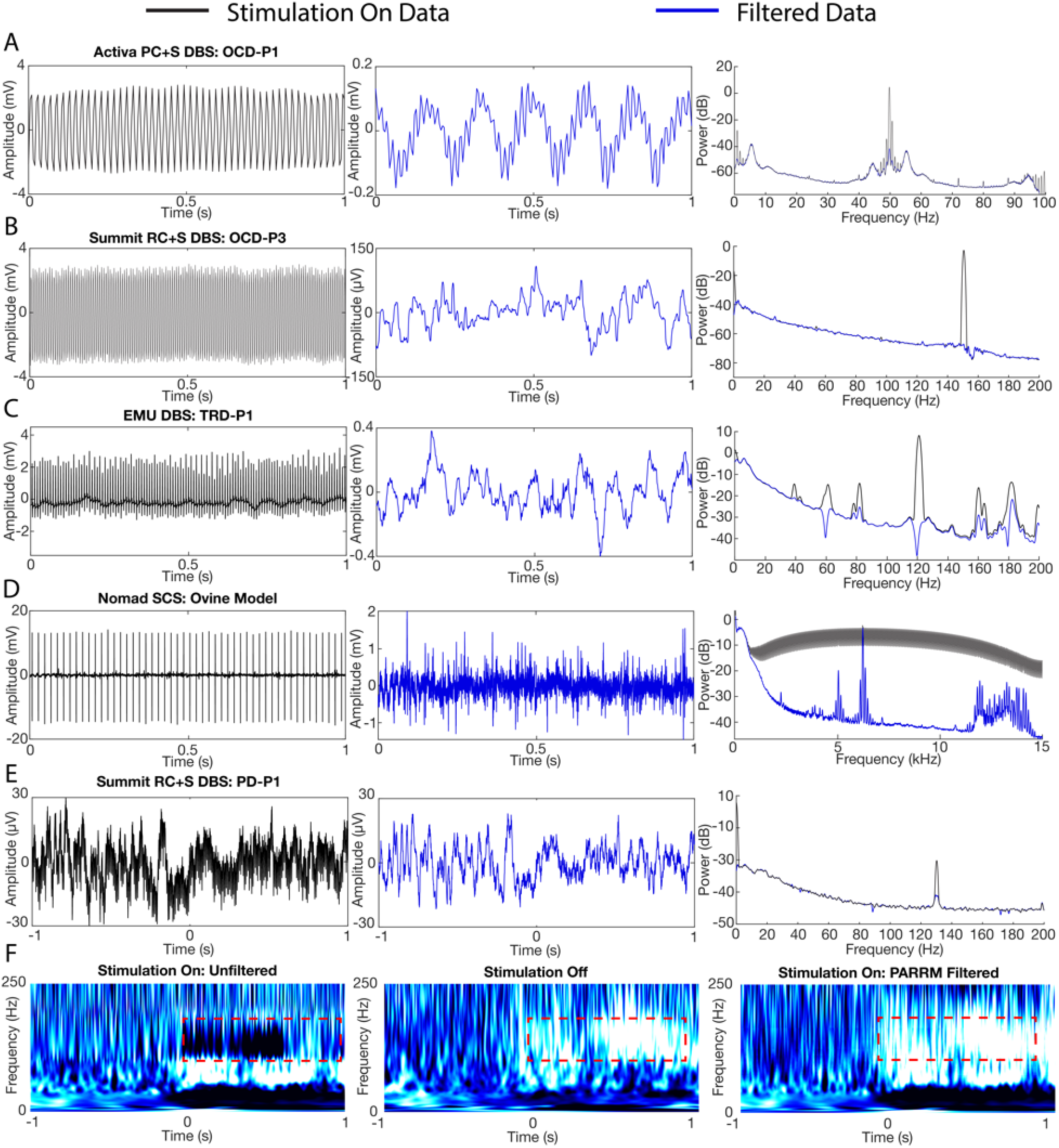
Demonstration of PARRM in human participants with DBS and SCS in ovine model. (a-d) Raw time-voltage LFP trace, PARRM filtered time-voltage LFP trace, and average PSD before (black) and after (blue) PARRM filtering, collected during (a) 150 Hz stimulation sampled at 200 Hz using Activa PC+S in OCD-P1 left VC/VS, (b) 150.6 Hz stimulation sampled at 1000 Hz using Summit RC+S in OCD-P3 right BNST, (c) 120 Hz stimulation sampled at 2000 Hz in TRD-P1 left ventral PFC during a cognitive control task, (d) 50 Hz spinal stimulation sampled at 30 kHz in ovine model using Ripple Nomad, (e) 130.2 Hz stimulation in STN sampled at 1000 Hz using Summit RC+S in PD-P1 recorded in right M1 during movement task. Left: one unfiltered trial in time domain. Center: PARRM filtered trial in time domain. Right: PSD of whole task (f) Averaged continuous wavelet transforms for a movement task zeroed to motion cue for stimulation on unfiltered data, stimulation off, and stimulation on PARRM filtered data recorded using the Summit RC+S in PD-P1 recorded in right M1. Location of high-gamma biomarker is indicated by the dashed red line.

### PARRM significantly attenuates stimulation artifacts from the Activa PC+S

We then applied PARRM to an extensive 1012 recording session dataset from two human neuropsychiatric DBS participants (NCT03457675). Prior to application of PARRM, the unfiltered electrophysiological signal recorded during stimulation for both participants displayed a large artifact, obscuring the LFP signal of interest (Figure 5a, left, S. Fig. 6a left). Following the application of PARRM, the amplitude of the resulting signal was reduced by a factor of 20. However, unexpected oscillations with non-stationary frequency content centered at approximately 6 Hz and 3 Hz for OCD-P1 and OCD-P2, respectively, remained after filtering (Figure 5a center, S. Fig. 6a center). Average power spectral densities were computed for all recordings and confirmed that the expected stimulation harmonics were well attenuated for both participants (Figure 5a right, S. Fig. 6a right).

**Figure 6:**
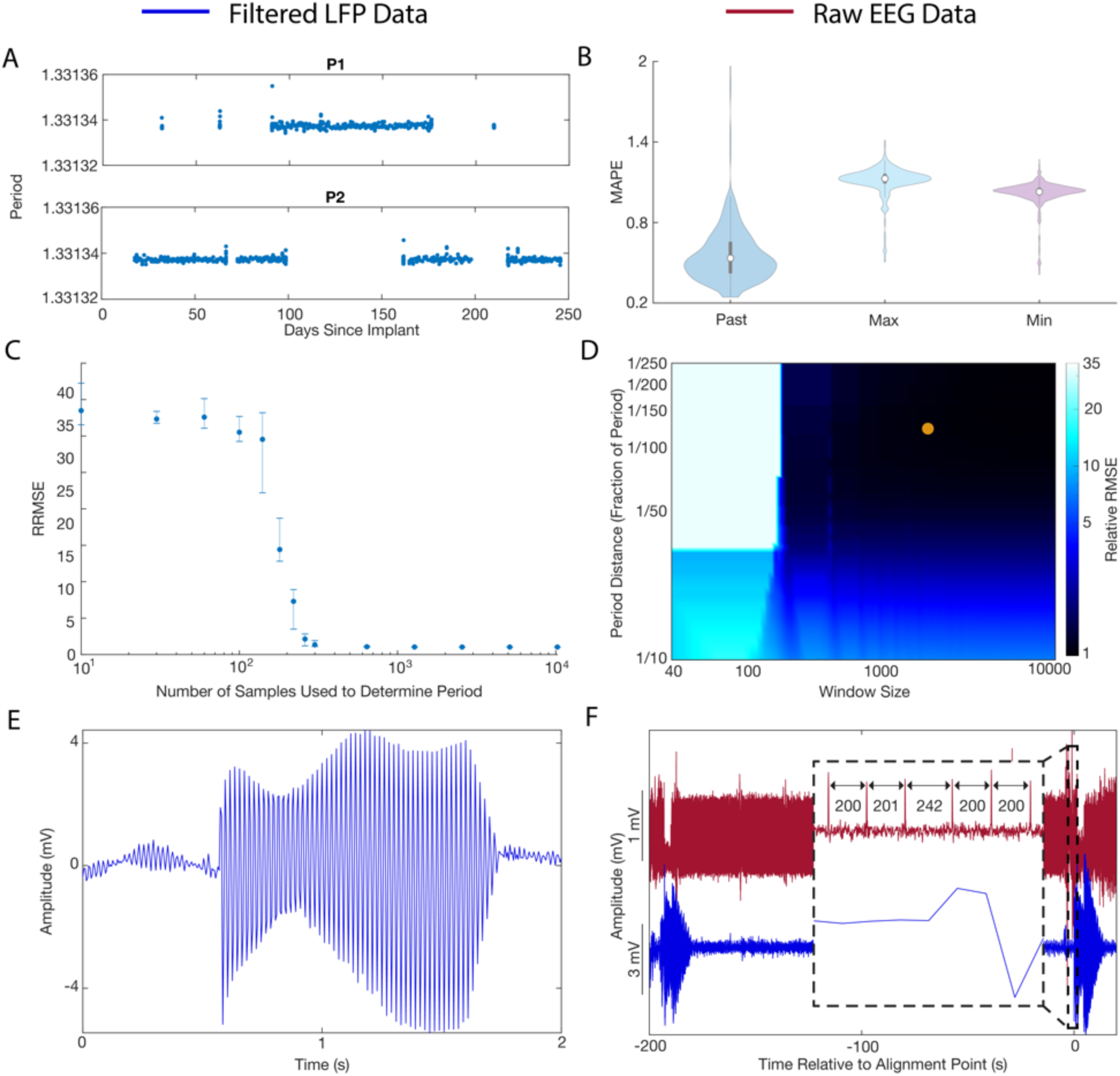
Practical considerations for implementing signal recovery via PARRM in real-time. (a) Exact period estimations in samples over 1012 recordings for P1 and P2 over 250 days since DBS implant.(b)Median absolute percent error (MAPE) between the standard PARRM filtering approach (using past and future samples, and exact period estimation) and using past samples only with an exact period estimation, using past samples only with the maximum period across the 1012 recordings, and past samples only with the minimum period across the 1012 recordings. (c) PARRM performance measured by relative root mean squared error (RRMSE) is dependent on the number of samples used to determine the period. Error bars show the spread (d) Heat map of RRMSE as a function of period distance (*D*_*period*_) and half window size (*N*_*bins*_). Darker blue indicates superior PARRM performance. Orange point indicates the *D*_*period*_ and *N*_*bins*_ that were used for all analysis. (e) Voltage-time LFP trace after PARRM filtering containing a jump in the period. (f) LFP (blue) and concurrent EEG (red), aligned using location of period jump identified in both recordings.

### PARRM removes artifacts in a wide range of therapeutic stimulation paradigms

After establishing the suitability of PARRM to deep brain recordings during DBS therapy, we evaluated the applicability of PARRM to different neuromodulation modalities to recover neural signals during 1) 150.6 Hz DBS using the Summit RC+S for OCD in the right BNST and left VC/VS (Figure 5b, S. Fig. 6b), 2) 120 Hz DBS in an EMU setting recording from left prefrontal cortex and amygdala (Figure 5c, S. Fig. 6c), 3) 50 Hz and 10 kHz SCS during rest in sheep recording approximately from spinal segments L5-S1 (Figure 5d, S. Fig. 6d), and 4) 130.2 Hz DBS using the Summit RC+S for PD in the STN recording from right M1 (Figure 5e). The effectiveness of PARRM for each setting was evaluated by comparing filtered recordings to unfiltered recording using raw time domain traces and power spectral densities. In all four modalities, PARRM was able to attenuate the stimulation artifacts at every amplitude and frequency leading to large reductions in artifact amplitude in both the time and frequency domains. PARRM was able to accurately remove artifacts and their harmonics at both low and high frequencies and, in the case of the EMU recording, identified and attenuated aliased artifacts. Lastly, PARRM was applied to data from M1 recorded using the Summit RC+S during a movement task where the subject was receiving concurrent 130.2 Hz DBS in the STN for PD. PARRM removed artifacts in the time domain on a single trial basis and reduced artifact amplitude in the frequency domain (Figure 5e). When all trials were averaged, PARRM was able to recover a known high-gamma biomarker for movement onset that was previously obscured by stimulation artifact (Figure 5f). Together, these analyses demonstrate that PARRM is readily adaptable to a wide range of neural recording paradigms and can enable the recovery of neural biomarkers otherwise obscured by stimulation artifacts.

### Potential for online application of PARRM in implantable technologies

The feasibility of implementing PARRM as an online method for low sampling rate recordings was investigated using LFPs sampled at 250 Hz by the Activa PC+S over 250 days. Using 1012 recordings from two Activa PC+S participants both in the clinic and at home, we estimated potential variability in the period over the 250-day span. Variation in the period was minimal (standard deviation of 10^−5^ samples) with the maximum and minimum differing from the median by only 2 × 10^−5^samples. The estimated period was consistent across both devices with only a 2 × 10^−7^sample difference between the medians (Figure 6a). Data filtered using past samples only and extreme periods were compared to data filtered using the standard PARRM approach where both past and future samples are used, and an accurate period is estimated. The similarity between these approaches was quantified using median absolute percentage error (MAPE). Using only past samples resulted in a MAPE of 0.6% when compared to a two-sided filter. Filtering using extreme periods and past samples resulted in a MAPE of approximately 1% when compared to a two-sided filter (Figure 6b). We then sought to estimate the minimum number of samples that were necessary to guarantee an accurate estimate for the period based on RRMSE with the simulated chirp data. RRMSE followed a roughly sigmoidal relationship with the number of samples used. RRMSE and its variability decreased for increasing number of samples. Improvement (decrease) beyond 1000 samples was minimal (1% difference) (Figure 6c). We then computed the RRMSE as a function of two filter parameters: the period distance (*D*_*period*_) and the half window size (*N*_*bins*,_). We found that increasing window size and decreasing period distance generally improved RRMSE. Improvement beyond a period distance of 1/150th of a period and 2000 samples was minimal (Figure 6d). Improvements in RRMSE were not fully explained by the total number of samples averaged for each parameter combination (S. Fig. 8). These results demonstrate that PARRM can be implemented with minimal onboard memory enabling real-time artifact removal.

### PARRM is robust to spurious changes in stimulation artifact

Lastly, we verified that PARRM is robust to spurious jumps in the stimulation period. After a jump in the period, PARRM temporarily filters using the incorrect phase of the artifact. However, due to the moving filter window, PARRM performance recovers a few seconds after a period jump (Figure 5e). These period jump events can be used to our benefit to align LFP recordings with external sensors, such as EEG (Figure 5f). These jumps can be located on the high sampling rate (30 kHz) EEG as increases in the difference between subsequent peaks. In the LFP, these events can be located by filtering using PARRM twice: once with a past window, and again with a future window. The peak in the product of the five-sample moving standard deviations of these two filtered recordings corresponds to the location of the alignment point (S. Fig. 9). These results demonstrate that PARRM can be robust to potential recording errors in an online environment and assist in temporal alignment of concurrent recordings.

## Discussion

In summary, we have developed a novel method, PARRM, that removes aliased electrical stimulation artifact in low and high sampling rate LFP recordings even in cases where the frequency of the underlying neural signal overlaps with the frequency of the aliased artifact (Figure 3). Before the development of PARRM, it was impossible to completely remove aliased artifacts resulting from stimulation frequencies or harmonics greater than the Nyquist frequency (125 Hz for recordings sampled at 250 Hz) without contaminating the underlying neural signal. Stimulation frequencies over 125 Hz are clinically relevant for PD, OCD, TRD, and pain (*16–18*). This capability opens the door for use of power efficient and high channel count implantable neurostimulation devices without sacrificing data quality, and is particularly useful for the two existing commercial DBS systems capable of concurrent stimulation and sensing at 250 Hz: the Percept, and the NeuroPace RNS (*19, 20*). PARRM is a low complexity algorithm that can develop templates for subtraction based on past data samples, requiring minimal computational resources and onboard storage, and could be implemented on existing and future neurostimulation devices.

Due to the conditions of our benchtop saline experiments, including impedance mismatch of electrodes in saline versus the human brain, most validation steps were completed via computational simulations. We chose to simulate our most limited recording scenario: the Activa PC+S at a sampling rate of 200 Hz. While the simulated waveform was not based on exact parameters of every component of the Activa PC+S device circuit, we believe that the simulations do not detract from the validation of the method. The simulated artifact waveform closely matches the reconstructed waveform observed during benchtop saline testing (S. Fig. 3). The simulation was an efficient way to evaluate PARRM performance over a vast array of DBS parameters and conditions. In the future, we hope that DBS device companies will consider publishing Simulink models of the DBS waveforms their devices produce to aid in artifact characterization and removal.

We found that when applying PARRM to Activa PC+S recordings, distinct, low-frequency, nonstationary oscillations remained. Nonstationary oscillatory artifacts, varying on a timescale shorter than the filter window, cannot be successfully mitigated using PARRM. We investigated whether these nonstationary oscillations may have been the product of variable noise, dependent on the phase of the waveform where a point was sampled, or, may have resulted from jitter in the true pulse location within a period compared to the pulse location predicted by PARRM. However, when this noise and jitter were incorporated into simulations, neither addition replicated the nonstationary oscillations (S. Fig. 7). Since these oscillations did not appear in saline recordings and could not be replicated via the addition of noise or jitter, we hypothesize that they arise from interactions between the electrical stimulation and the unique chemical medium and structural environment of the brain and should be investigated further in future studies. Recording configurations which minimize such artifacts, as well as aperiodic artifacts from other sources, are valuable for maximizing PARRM performance.

Online, real-time artifact removal via PARRM will enable unbiased exploration of neural biomarkers that may have previously been obscured by stimulation artifact. More broadly, PARRM may be applicable in any domain in which a periodic artifact should be removed to recover an underlying signal of interest. While more development is required before PARRM can be applied to do onboard artifact rejection during concurrent neurostimulation therapy and sensing, PARRM could ultimately contribute to the accurate detection of neural biomarkers and the development of closed-loop neuromodulation therapies.

## Materials and Methods

### I. Human LFP recordings from implanted DBS devices for OCD

Research subjects were four participants, each with a history of long-standing OCD, that underwent clinically indicated DBS surgery for treatment of OCD. Two participants (OCD-P1 and OCD-P2) were implanted with the Activa PC+S (Medtronic, Minneapolis, MN, USA) device, and two participants (OCD-P3 and OCD-P4) were implanted with the Summit RC+S (Medtronic, Minneapolis, MN, USA) device. Each participant gave fully informed consent according to study sponsor guidelines, and all procedures were approved by the local institutional review board at Baylor College of Medicine (H-40255, H-44941).

DBS leads (Model 3778) were intracranially placed bilaterally in the VC/VS or BNST based on clinical indications and connected to the Activa PC+S or Summit RC+S to enable control of DBS and LFP recordings. OCD-P1 received bilateral stimulation while OCD-P2 received unilateral stimulation. LFP was sensed with bipolar contacts around the stimulation contact at a sampling rate of 200 Hz (Activa PC+S) or 1000 Hz (Summit RC+S). Scalp EEG sampled at 30 kHz was concurrently recorded using tripolar concentric ring electrodes (tCRE, CRE-Medical, University of Rhode Island, RI, USA).

### II. Intracranial electroencephalography recordings

A research subject with a history of treatment-resistant depression (TRD-P1) was implanted with clinical depth electrodes (PMT, Chanhassen, MN, USA) spanning the amygdala, prefrontal cortex, orbitofrontal cortex and cingulate cortex, as well as bilateral DBS electrodes (Vercise Gevia; Boston Scientific, Marlborough, MA, USA) in the VC/VS and subcallosal cingulate. Research protocols were approved by the institutional review board at Baylor College of Medicine (H-43036, H-40255), and the research subject provided written and verbal voluntary consent to participate in the study.

Intracranial electroencephalographic (iEEG) signals from depth electrodes were recorded at 2 kHz with a bandpass of 0.3-250 (4th order Butterworth filter) using a 256 channel Blackrock Cerebus system (Blackrock Microsystems, Salt Lake City, UT, USA). Stimulation was concurrently delivered through DBS electrodes using Cerestim (Blackrock Microsystems, Salt Lake City, UT, USA) to deliver continuous stimulation at 130 Hz, 100 µS pulse width and 4-6 mA. In order to remove line noise, notch filters were applied at 60, 120, and 180 Hz.

### III. Sheep spinal electrophysiological recordings

One sheep underwent surgery to implant a custom-built 24 contact SCS device on the epidural surface of the spinal cord from approximately the L5-S1 spinal segments. All study procedures were conducted with the approval of the Brown University Institutional Animal Care and Use Committee (19-04-0002) and in accordance with the National Institutes of Health Guidelines for Animal Research (Guide for the Care and Use of Laboratory Animals). Device wires were externalized and connected to a Nomad (Ripple Neuro, Salt Lake City, UT, USA) neural interface system to allow for simultaneous stimulation and recording of the spinal cord at 30 kHz. Stimulation was controlled by a custom-written MATLAB (Mathworks, Natick, MA, USA) script to deliver current at levels typically used for chronic pain management using SCS (0-2000 μA, 50 Hz and 10 kHz).

### IV. Human LFP recordings from implanted DBS devices for PD

One PD patient (PD-P1) was implanted with bilateral cylindrical quadripolar deep brain stimulator leads into the subthalamic nucleus (STN, Medtronic model 3389) and bilateral placement of paddle-type quadripolar cortical paddles into the subdural space over motor cortex (MC, Medtronic model 0913025). Each pair of STN and MC leads was connected bilaterally to a Summit RC+S device in a pocket over the pectoralis muscle (Medtronic Summit RC+S model B35300R). The paddle lead was placed in the subdural space through the same frontal burr hole used for the subthalamic lead. At least one contact covered the posterior precentral gyrus (presumed primary motor cortex), approximately 3 cm from the midline on the medial aspect of the hand knob. The STN leads was implanted in the motor territory of the STN. Placement was confirmed with movement-related single-cell discharge patterns. The study was approved by the hospital institutional review board (IRB) at University of California San Francisco Medical Center under a physician sponsored investigational device exemption (G180097) and was registered at ClinicalTrials.gov (NCT03582891). The patient provided written consent in accordance with the IRB and Declaration of Helsinki.

### V. Period-based Artifact Reconstruction and Removal Method (PARRM)

At each time bin *t*, PARRM subtracts an estimate of the stimulation artifact at time bin *t* from the recorded signal at time bin *t* (Figure 2). The estimate of the stimulation artifact is formed by averaging the recorded signal at other time bins that are in a temporal region near time bin *t* and also approximately at the same phase of stimulation as time bin *t*. The artifact is presumed to be roughly identical for all of these time bins, including time bin *t*. Averaging reduces the influence of brain signals and additional sources of noise, so that the subtracted signal is primarily artifact.

Let *T* denote the stimulation period relative to the sampling rate (in units of sampling time bins). The time bins included in the average are those times bins *s* such that

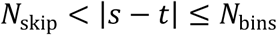

and such that

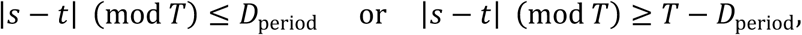

where a (mod *T*) denotes a modulo *T*, and where 0 ≤ *N*_skip_ < *N*_bins_ and 0 ≤ *D*_period_ ≤ *T* are user-chosen design parameters. (The additional criterion *s* − *t* < 0 can be included so that only past observations are used to estimate the stimulation artifact; see Supplementary Materials.) Let B_t_ denote the collection of those times bins *s* that are used for averaging and let |*B*_*t*_| denote the number of such time bins. Using *r*_*t*_ to denote the recorded signal at time bin *t*, the corrected signal is *c*_*t*_ defined by

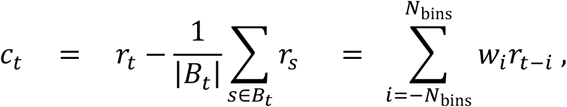

where *w*_*i*_ is a list of weights defined by *w*_0_ = 1, and *w*_*i*_ = −1/|*B*_0_| if −*i* ∈ *B*_0_, and *w*_0_ = 0 otherwise. The final expression shows that the PARRM correction can be implemented by a fixed linear filter (with the filter weights denoted by *w*_*i*_), making it fast and simple to implement. (If the additional criterion *s* − *t* < 0 is used, then the final summation would begin at *i* = 0. The final formula also needs to be modified near the start and end of stimulation; see Supplementary Materials.)

The design parameters for the PARRM filter are *N*_bins_, *N*_skip_, and *D*_period_. Larger choices of *N*_bins_ allow more data to be averaged in order to estimate the artifact, reducing estimation variability. But, larger choices of *N*_bins_ also lengthen the temporal window used to estimate the artifact, perhaps introducing estimation bias if the artifact shape is changing in time. Because neural signals have temporal autocorrelation, it is important to avoid averaging data too close to time bin *t* or the neural signal itself could be subtracted during artifact removal. Larger choices of *N*_skip_ help to mitigate this danger, but also reduce the amount of data used to estimate the artifact. Similar to *N*_bins_, larger choices of *D*_period_ allow more data to be averaged, but also introduce more estimation bias by temporally smoothing the estimated artifact. The optimal choices for these design parameters will vary depending on the situation; see Supplementary Materials.

### VI. Period estimation

PARRM needs a precise estimate *T*of the stimulation period relative to the sampling rate. *T* can be determined via several methods. This paper uses an automated, data-driven method that works by searching for a period that creates a strongly resolved template (Fig. 2E, S. Fig. 1). For each candidate period *δ* > 0, the method estimates a waveform template with this period and then quantifies deviation from the estimated template. The candidate period with the smallest deviation is selected as the final estimate *T*of the period that is used by PARRM. A similar period finding method was described by Tzou et. al (*21*).

Let *m* ≥ 0 be an integer. For each potential period *δ* > 0 and each parameter vector *β* = (*β*_1=_, …, *β*_2m+1_) define the functions

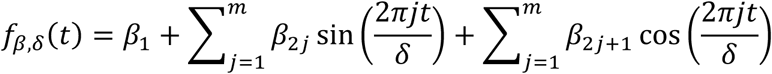

The function *f*_*βδ*_ is a periodic function with period *δ*. Each *f*_*βδ*_is a candidate artifact waveform. The parameter vector *β* controls the strength of the different frequencies that define *f*_*βδ*_, and *m* controls the number of allowed frequencies. Let Q((*t*_*K*_, *y*_*K*_): *k* = 1, …, *n*)V be a collection of (time, value) pairs. The *y*_k_ value used here is the change in recorded LFP amplitude at time *t*_k_ with some preprocessing to obtain standardized units, reduce the influence of outliers, and reduce the size of the dataset; see Supplementary Materials. Mean squared error is used to measure how well the function *f*_*βδ*_ fits these pairs:

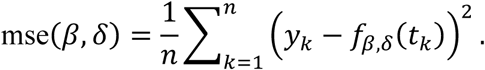

For fixed *δ*, the optimal *β*, say, 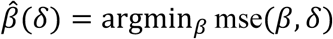, can be computed exactly using linear regression techniques; see Supplementary Materials. The final estimate of the period is

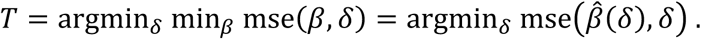

The minimization over *δ* is complicated by many local minima, spurious ‘distractor’ solutions that mimic the harmonics of the true waveform (S. Figure 1), and high sensitivity to small changes in*δ*. The examples in this paper use a penalized, stagewise search that begins with smaller intervals of data (to reduce the sensitivity to *δ*), smaller *m* (to reduce the number of local minima), and a penalty for higher frequency solutions (to help avoid distractor solutions); see Supplementary Materials. This seems to be the most delicate part of the period-finding procedure. Once *T* is found, it is fixed for PARRM. Simpler methods for period finding are under active development and will be described in a future publication.

### VII. Implementation of state-of-the-art filters

Hampel filter, moving average subtraction, matched filter, and notch filter performance were used as a comparison point to PARRM performance. Hampel filters interpolate artifactual components in the frequency domain and have been shown to be an effective approach for removing DBS artifacts in EEG recordings. We implemented a standard Hampel filter in MATLAB based on the method described by Allen et al. (*22*). Moving Average Subtraction (MAS) employs peak finding to identify each stimulation pulse in an up-sampled recording before averaging neighboring pulses to construct a local template, and has been shown to be effective in signal recovery for low and high sampling rate EEG recordings during DBS. Following the method described by Sun and Hinrichs et al., we implemented a similar filter in MATLAB (*23*). Matched filters estimate the amplitude and phase of a series of sinusoidal harmonics of the artifact by maximizing cross correlation and have been shown to be effective in signal recovery for simulated DBS artifacts added to EEG data. We implemented a matched filter in MATLAB using six matched components based on the method described by Sun et al. (*24*). Notch filters at the stimulation frequency and its harmonics are an effective method for removing DBS artifacts by completely attenuating power at affected frequencies. Second order Infinite Impulse Response (IIR) notch filters with a half-power distance of 5 Hz were applied at the stimulation harmonics and their aliases using the MATLAB *designfilt* and *filtfilt* functions. For recordings sampled at 200 Hz, a high-pass finite impulse response (FIR) filter with 2 dB stopband attenuation, transition band between 2 and 3 Hz, and a passband ripple of 0.1 dB and a 20th order low-pass IIR filter at 97 Hz with 0.1 dB of passband ripple were used to attenuate the aliased components at 0 and 100 Hz (for 150 Hz stimulation).

### VIII. Experimental validation of PARRM in saline

The artifact removal method was validated by simulating the recording conditions in the brain using a setup in a saline solution. The DBS lead (Model 3778) and case were immersed on opposite sides of a plastic container containing 1x phosphate buffered saline solution at room temperature. A platinum electrode connected to a waveform generator was placed adjacent to the stimulating electrode to simulate LFP (S. Fig. 2). Single frequency (10 Hz and 50 Hz) oscillations were injected by the waveform generator alongside 2 V, 150 Hz, 90 µs pulse width stimulation. The efficacy of the removal method was characterized by comparing the distributions of absolute errors of the artifact free injected signal with unfiltered, moving average subtraction (MAS) filtered, notch filtered, and PARRM filtered signals. Baseline noise was estimated during recordings where stimulation was off and there was no injected signal from the waveform generator. Significant differences from baseline noise were estimated using a Wilcoxon-Rank Sum test.

### IX. Experimental validation of PARRM using Simulink

The recording circuit for the Activa PC+S device was simulated using Simulink (Mathworks, Natick, MA, USA) (*25*) (S. Fig. 3A). The simulation input was a train of modeled DBS pulses sampled at 120 kHz, and the output was the simulated stimulation waveform as if it were being recorded by the Activa PC+S (S. Fig. 3B). The Simulink model is publicly available on GitHub (see Availability statement). By default, a stimulation frequency of 150 Hz, amplitude of 2 V, and pulse width of 90 µs were used. The simulation reached a steady state after two seconds. The final stimulation waveform was then used to create pulse trains that match the simulated injected signal in length. Each simulated pulse train was downsampled by a factor of 601 or 121 (199.67 Hz or 992 Hz) to replicate the true sampling rate, which deviates slightly from the sampling rate stated by the device (200 Hz or 1000 Hz). For each simulation, the stimulation pulse train was added to a series of 30 linear chirps. Each chirp was two seconds in length and separated from the following chirp by one second with 0.1 seconds of jitter. Chirp amplitude was twice the root mean squared amplitude of the baseline noise. Gaussian noise equal in magnitude to what was observed in saline was added to each simulation. For the signals sampled at roughly 200 Hz, chirps ranged from 0 to 100 Hz. For the signals sampled at roughly 1000 Hz, chirps ranged from 0 to 200 Hz. PARRM performance using simulated data was compared to that of a hampel filter, MAS filter, matched filter, and notch filter. A parameter sweep was run to test PARRM performance across varying stimulation frequencies (80-180 Hz), amplitudes (0.5-5 V), pulse widths (30-180 µs), and chirp lengths (1-10 s).

### X. Spectral analysis

Time frequency decomposition was performed using a continuous complex Morlet wavelet transform. For data sampled at 200 Hz, 500 steps from 0 to 100 Hz were used. Wavelets were constructed using one cycle at the minimum frequency up to 20 at the maximum frequency. Steps were linearly spaced for analysis of chirp signals and logarithmically spaced for analysis of stationary sinusoidal signals. For data sampled at 1000 Hz, 500 linearly spaced steps from 0 to 200 Hz were used. Wavelets were constructed using one cycle at the minimum frequency up to 30 at the maximum frequency also with linearly spaced steps. For analyzing the frequency content of each chirp, we computed a windowed power spectral density using the decomposition. The power for each frequency was computed by averaging the power in a window centered at the time the frequency of interest occurred during the linear chirp. The window size was four samples for the 200 Hz recordings and 20 samples for the 1000 Hz recordings. Stationary power spectral densities were computed using the MATLAB *pspectrum* function.

### XI. Estimation of filter performance

#### 1. Visual comparison: averaged chirp

In order to visually compare the different filtering approaches, all 30 chirps were averaged together to produce a single average chirp. This method was used to visually show how well each filter was able to recover the signal over many trials.

#### 2. Frequency domain chirp comparison metric: Windowed PSD

In order to compare how well each filtering approach was able to recover the chirp signal in the frequency domain, the distribution of power was compared for each frequency. Power was computed by calculating the decibel ratio of the signal of interest and the concurrent noise. Significant differences from the artifact free signal (chirp without simulated DBS) at each frequency were computed using a 2-sample t-test.

#### 3. Time domain chirp comparison metric: Relative root mean squared error

In order to compare how well each filtering approach was able to recover the chirp signal in the time domain, the distributions of relative root mean squared error were compared. Relative root mean square error (RRMSE) was calculated for each chirp by dividing the root mean squared error between the filtered and theoretical chirp signals by the root mean squared error of the artifact free and theoretical chirp signals.

#### 4. Parameter sweep metric: Relative R Ratio

In order to compare how well each filtering approach was able to recover the chirp signal in the frequency domain as a whole, the distribution of relative R ratios was computed (*26*). Relative R ratio was computed as

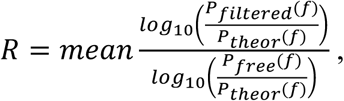

where *P*_*filtered*_ is the power for the filtered signal, *P*_*theor*_ is the power for the theoretical chirp signal (without noise), and *P*_*free*_ is the power for the chirp signal without stimulation artifact.

### XII. Movement Task

A movement task written using jsPsych was presented to PD-P1 on a laptop touch screen computer (*27*). The patient was presented with a target appearing in one of four locations on the screen followed by a cue to move and a baseline period (each lasting up to three seconds). The patient performed 60 reaches (15 to each target, randomized) with therapeutic deep brain stimulation off or on in the STN. Synchronization of neural data and task data was done using the clock of the patients’ study computer. Two channels were recorded from motor cortex with a 1000 Hz sampling rate.

For movement-related changes in spectral power, data were filtered using a two way 3rd order FIR filter (eegfilt from eeglab toolbox with fir1 parameters) and bandpassed in frequencies between 1-200 Hz (*28*). Data from all trials were aligned relative to the onset of movement and averaged. The averaged amplitude was normalized by a 1000 ms window prior to cue presentation (time 0). Data were z-scored by subtracting the average baseline amplitude and dividing by the baseline standard deviation. This z-score procedure was performed for each frequency separately.

### XIII. Feasibility for use of PARRM as an online method

Using 1012 recordings from two human participants implanted with the Activa PC+S, we investigated whether it would be feasible to implement PARRM using an existing device. For PARRM to be effective as an online method, filter performance should depend on past samples only (rather than past and future samples) and should be robust to any foreseeable variation in the stimulation period over time. Additionally, the recording duration required to make an initial period estimate should be minimal, and filtering should require minimal resources onboard the device. To this end, a 40-second-long segment from each of the 1012 recordings was filtered using PARRM. The period was estimated for each recording. Extreme periods were identified by finding the maximum and minimum period over all the 1012 recordings. Data filtered using past samples only and extreme periods were compared to data filtered using the previously described approach where both past and future samples are used, and an accurate period is estimated. In total, the data were filtered using (1) past and future samples and accurate periods, (2) past samples only using accurate periods, (3) past samples only using minimum extreme periods, and (4) past samples only using maximum extreme periods. In order to quantify the magnitude of difference between the four filtering approaches, the median absolute percentage error (MAPE) between the original approach and the alternative approach was computed for each recording. Additionally, the RRMSE was found as a function of the number of samples used to determine the period, the period distance, and the window size for simulated chirps sampled at 200 Hz.

### XIV. LFP synchronization with external sensors

For one of the human participants (P2) implanted with the Activa PC+S, we synchronized the LFP recording with concurrent EEG. Synchronization was achieved by identifying ‘jumps’ in the stimulation period which occurred simultaneously in both recordings. Jumps in the difference between EEG peak times found using the MATLAB *findpeaks* function were used to locate these events in the EEG. In LFP recordings, these events were located by comparing data filtered using only past versus only future samples. A moving standard deviation with a window of five samples was computed for both recordings and the ‘jump’ corresponded to the peak in their product.

## Acknowledgements

We thank the participants and their families for participating in this research. We also thank Kendall Lane for illustrating Figure 1. Activa PC+S and Summit RC+S devices were donated by Medtronic.

## Author contributions

MTH, EDVR, NRP, and DAB conceived of the Period-based Artifact Removal Method. EDVR, JSC, and RG carried out all data analysis with input from NRP, MTH, and DAB. NRP and EDVR conducted saline validation experiments. NRP, EM, GSV, MAO, ACV, NR, and WKG conducted LFP recordings with OCD participants. JSC, RD, and SS conducted ovine SCS experiments. ABA, DNO, KRB, and SAS conducted iEEG recordings with TRD participants. RG, RW, and PAS conducted LFP recordings during the movement task with PD participants. NRP, EDVR, JSC, RG, ABA, MTH, and DAB wrote the manuscript. DAB and MTH oversaw manuscript conception and completion. All authors reviewed and edited the manuscript.

## Data and materials availability

Data and code used to produce this manuscript is available on GitHub (https://github.com/neuromotion/PARRM).

## Supplemental Figures

### Materials and Methods

**S. Fig. 1**. Distractor period mimics a harmonic of the true DBS waveform.

**S. Fig. 2**. Experimental saline setup.

**S. Fig. 3**. Simulated Activa PC+S frontend filtering circuit and output DBS waveform.

**S. Fig 4**. Continuous wavelet transforms of simulated chirps

**S. Fig 5**. Simulations show that PARRM is effective at a wide range of DBS parameters.

**S. Fig 6**. Additional demonstration of PARRM in human participants with DBS, iEEG recordings during concurrent DBS, and Spinal Cord Stimulation in ovine model.

**S. Fig. 7**. Exploration of non-stationary oscillations leftover after PARRM in human data.

**S. Fig. 8**. Number of samples averaged as a function of window size and period distance.

**S. Fig. 9**. Illustration of method for finding period jumps in LFP.

