## Supplemental text for "Uncovering biomarkers during therapeutic neuromodulation with PARRM: Period-based Artifact Reconstruction and Removal Method"

### Materials and Methods

I. PARRM derivation and modifications

Here we give details of the derivation that PARRM can be implemented as a linear filter. The corrected signal is defined in the main text (Methods) as

$$c_t \quad = \quad r_t - \frac{1}{|B_t|}\sum_{s \in B_t} r_s \, .$$

Ignoring edge effects and inspecting the definition of $B_t$, we see that $s \in B_t$ if and only if $s - t \in B_0$. Or, equivalently, $u \in B_0$ if and only if $u + t \in B_t$. Hence, we can express

$$c_t \quad = \quad r_t - \frac{1}{|B_0|}\sum_{u \in B_0} r_{u+t} \quad = \quad r_t - \frac{1}{|B_0|}\sum_{i=-N_{\text{bins}}}^{N_{\text{bins}}} \chi_{B_0}(i) r_{t+i}$$

$$= \quad r_t - \frac{1}{|B_0|}\sum_{i=-N_{\text{bins}}}^{N_{\text{bins}}} \chi_{B_0}(-i) r_{t-i} \, ,$$

where $\chi_B(a) = 1$ if $a \in B$ and $\chi_B(a) = 0$ if $a \notin B$. Defining the weight vector $w$ as in the main text and observing that $\chi_{B_0}(0) = 0$, we see that the final expression can be expressed as the convolution

$$c_t = \sum_{i=-N_{\text{bins}}}^{N_{\text{bins}}} w_i r_{t-i} ,$$

which is the familiar formula for a discrete linear filter. Note that the derivation is unchanged if we modify $B_t$ to have the additional restriction that $s - t < 0$, meaning that only past observations are used to estimate the stimulation artifact.

This derivation ignores edge effects that occur for those $t$ such that $t - N_{\text{bins}}$ or $t + N_{\text{bins}}$ extends beyond the recording period or is outside of the period of stimulation. We use the original definition of $c_t$ in these cases, where $B_t$ includes only those time bins that are recorded during the stimulation period.

### II. Period finding

The main text (Methods) describes the optimization criterion for selecting the stimulation period $T$ used by PARRM. Here we describe the exact preprocessing used to create the $y_k$ values, the details of linear regression optimization step, the details of the stagewise search over candidate periods $\delta$, and the values of the design parameters.

*Preprocessing.* We begin with an electrophysiological signal recorded during stimulation. Each channel of the recording is processed as follows: (1) The signal is differenced, i.e., the signal at time step $t - 1$ is subtracted from the signal at time step $t$. Differencing helps remove low frequency modulations unrelated to the artifact. (2) The (differenced) signal is divided by the mean absolute value of the (differenced) signal to give a unitless, standardized measurement. (3) Any values of this standardized signal greater than 3 are set to 3 and any values less than -3 are set to -3. This clipping of large magnitude observations helps remove the influence of outliers. The preprocessing results in a collection of $(t_k, y_k)$ pairs, where $t_k$ is the time bin and $y_k$ is the final (differenced, standardized, clipped) signal. Finally, we subsample these pairs in some way to obtain a smaller dataset, e.g., by taking all pairs from a shorter interval or by choosing pairs uniformly at random (without replacement). This subsample is the collection of $\big((t_k, y_k): k = 1, \dots, n\big)$ pairs referenced in the main text (Methods) and below.

*Linear regression*. For each $(t_k, y_k)$ pair, define the row vector $x_k = (x_{k1}, x_{k2}, \dots, x_{kM})$ for $M = 2m + 1$ by $x_1 = 1$, $x_{2j} = \sin(2\pi j t_k/\delta)$, and $x_{2j+1} = \cos(2\pi j t_k/\delta)$, for $j = 1, \dots, m$. Then

$$\mathrm{mse}(\beta, \delta) = \frac{1}{n}\sum_{k=1}^{n}\left(y_k - f_{\beta,\delta}(t_k)\right)^2 = \frac{1}{n}\sum_{k=1}^{n}\left(y_k - \beta \cdot x_k\right)^2,$$

where $a \cdot b$ is the usual Euclidean dot product. This is the usual mean squared error formulation of simple linear regression. The minimizing $\beta$ is well known to be

$$\hat{\beta}(\delta) = \mathrm{argmin}_\beta\, \mathrm{mse}(\beta, \delta) = (x^T x)^{-1} x^T y\,,$$

where $x$ is the $n \times M$ matrix with $k$th row $x_k$, where $y = (y_1, \dots, y_n)$ is a column vector, and where the final expression uses standard notation for matrix transpose, inverse, and multiplication.

*Period search.* As described in the main text (Methods) we want to compute

$$T = \mathrm{argmin}_\delta\, \mathrm{mse}\big(\hat{\beta}(\delta), \delta\big)$$

or, if we have $L$ simultaneously recorded channels, then we want to compute

$$T = \mathrm{argmin}_\delta \sum_{\ell=1}^{L} \mathrm{mse}\big(\widehat{\beta_\ell}(\delta), \delta\big)\,,$$

where $\widehat{\beta_\ell}(\delta)$ is the linear regression solution for the $\ell$th channel. We use Matlab's `fminsearch` function (with the default parameters) which is an implementation of the Nelder-Mead simplex method as described in Lagarias, 1998 *(29)*. The search requires an initial guess $\delta_0$. We use a complicated method to arrive at a good initial $\delta_0$.

Define the function

$$h(\beta) = \sum_{j=1}^{2m+1} p_j \beta_j^2$$

for $p_j = j/(2m^2 + 3m + 1)$, which makes $p$ sum to one. The function $h$ serves as a penalty to express a preference for $\beta$'s that emphasize low frequencies, helping to avoid higher frequency 'distractor' solutions that mimic the harmonics of the true waveform (S. Figure 1). Now define the function

$$F(\delta, \lambda) = \mathrm{mse}(\hat{\beta}(\delta), \delta) + \lambda h(\hat{\beta}(\delta))$$

or, for multiple channels,

$$F(\delta, \lambda) = \sum_{\ell=1}^{L} \mathrm{mse}\big(\widehat{\beta_\ell}(\delta), \delta\big) + \lambda h(\widehat{\beta_\ell}(\delta))\,.$$

An initial estimate for the stimulation period is made by dividing the sampling rate by the DBS stimulation frequency (e.g. 1.33 = 200 Hz sampling rate / 150 Hz stimulation rate). A grid of 201 points with a spacing of $10^{-4}$ and a grid of 201 points with a spacing of $10^{-5}$ are centered at this initial estimate. For each of these grid points $\delta$, we evaluate $F(\delta, 1)$ using $m = 5$ and the $n = 5000$ data points at the center of the recording. We select the five $\delta$ that give the lowest values of $F(\delta, 1)$ and use these as five separate initialization points for Matlab's `fminsearch` function to minimize $F(\delta, 1)$. Of the five optimized $\delta$, we choose the one with smallest $F(\delta, 1)$. This $\delta$ becomes the new seed for a second stage of the search. The second stage is like the first, except that we build the grids around the new seed $\delta$. Also, the grids have spacings of $10^{-4}/2$ and $10^{-5}/2$, respectively, and we use $m = 10$ and $n = 10000$. The optimal $\delta$ found in the second stage is used to seed the third and final stage. Now the grids have spacings $10^{-4}/3$ and $10^{-5}/3$, respectively, and we use $m = 20$ and $n = 25000$. Also, the $n = 25000$ data points are chosen uniformly at random (without replacement) from the recording. The final $\delta$ selected from this third stage is used as the initial starting point $\delta_0$ of the search described in the main text to obtain

$$T = \text{argmin}_{\delta} F(\delta, 0)$$

with $m = 20$ and with the same $n = 25000$ datapoints from stage 3 of the search. Note that there is no penalty for spurious solutions in the final optimization.

We developed this search procedure in an ad hoc manner experimenting with several recordings of human LFP using the PC+S device and then found that it worked well across a variety of datasets and stimulation devices. Many of the choices and parameters are arbitrary and we have not carefully explored the effects of changing them. Visual inspection of the windowed data and how tightly it adheres to the resulting waveform makes it easy to verify if the procedure has worked properly. We are actively developing simpler and more robust period finding approaches that integrate more tightly with the PARRM filtering, but they will be described in future work.

### III. Filter parameter selection

The first parameter chosen when optimizing PARRM for a new recording was the period distance $D_{period}$. We decided on the value of $D_{period}$ depending on the shape of the reconstructed stimulation waveform. We identified the size of the smallest feature in the waveform in samples and chose $D_{period}$ to be small enough such that samples within that feature would be relatively stationary. For the PC+S, RC+S, SCS, and EMU recordings this parameter was set to 0.01, 0.01, 0.005, 0.005 samples respectively. We then increased the half window size $N_{bins}$ until the artifact appeared to be significantly attenuated in both the time and frequency domains. For the PC+S, RC+S, SCS, and EMU recordings the final value of this parameter was set to 2000, 6000, 6000, 1000 samples respectively. Choice of the value for $N_{skips}$ would become more relevant once a signal of interest was known so as to avoid including samples within that signal. This parameter was set to 20 samples for all recording paradigms. In the future, we intend to automate the choice of these parameters.

IV. Exploration of non-stationary oscillations leftover after PARRM in human Activa PC+S data

Given the presence of residual nonstationary oscillations following filtering using PARRM, we investigated whether they were sourced froms features we initially considered to be negligible; namely noise dependent on the artifact phase and jitter in the artifact peak location (S. Fig. 7). In order to find the phase dependent noise, we overlaid all samples on the time scale of one period. A 1000 sample moving standard deviation was then computed to calculate the expected noise as a function of artifact phase. Gaussian noise with standard deviation corresponding to the phase of each sample was then added to LFP artifacts reconstructed using PARRM in order to evaluate whether the noise recreated the nonstationary oscillations that PARRM was unable to remove. Since we concurrently recorded high resolution (30 kHz) EEG, we were able to estimate the true locations of the artifact pulses using this recording. In order to quantify any jitter in the location of the EEG artifact peaks, we reconstructed a typical EEG artifact using PARRM. We then convolved this reconstruction with EEG upsampled by a factor of 10 using linear interpolation in order to estimate the jitter in peak location for each artifact. This jitter was then added to LFP artifacts reconstructed using PARRM in order to evaluate whether the addition recreated the nonstationary oscillations that PARRM was unable to remove.

V. Utilization of PARRM for alignment of LFP to externalized sensors

Through the use of a moving filter window, PARRM performance is able to recover in the presence of changes to the stimulation waveform such as a jump in the period present prior to stimulation off in Activa PC+S recordings (Figure 6e). Using such events, the same foundational principles of the artifact removal method can be used to align LFP to external sensors with high precision. Jumps in stimulation period reliably occur during Activa PC+S recordings seconds before stimulation turns off. The location of this period jump event can be precisely identified on high sampling rate external sensors, such as EEG, via peak finding, and located on LFP via forward and backward PARRM filtering (Figure 6f, S. Fig. 9). The location of the jump in period can then be used as a synchronization point across both recordings. Reliable changes in stimulation waveform could be used in tandem with PARRM to synchronize data from other neurostimulation devices with external sensors. External sensors such as electrocardiogram or video can be used to record moment-to-moment changes in behavior or symptoms. This behavioral information can then be correlated with LFP activity to identify neural signatures of relevant behavioral states. Forwards and backwards filtering using PARRM enables the time synchronization of LFP recordings with recordings from external sensors that is essential for biomarker identification.

**Supplemental Figures:**

**S. Fig. 1.** Distractor period mimics a harmonic of the true DBS waveform.

**S. Fig. 2.** Experimental saline setup.

**S. Fig. 3.** Simulated Activa PC+S output DBS waveform.

**S. Fig 4.** Continuous wavelet transforms of simulated chirps

**S. Fig 5.** Simulations show that PARRM is effective at a wide range of DBS parameters.

**S. Fig 6.** Additional Demonstration of PARRM in human participants with DBS, iEEG recordings during concurrent DBS, and Spinal Cord Stimulation in ovine model.

**S. Fig. 7.** Exploration of non-stationary oscillations leftover after PARRM in human data.

**S. Fig. 8.** Number of samples averaged as a function of window size and period distance.

**S. Fig. 9.** Illustration of method for finding period jumps in LFP.

A

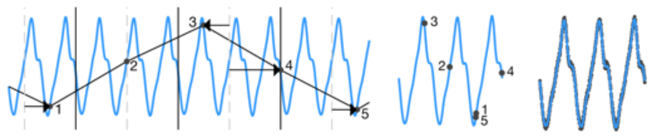

**Supplementary Figure 1: Distractor period mimics a harmonic of the true DBS waveform.** (a) Light blue trace indicates reconstruction of the estimated template waveform. Black trace indicates raw LFP sampled at 200 Hz. Black points indicate individual raw LFP samples. The distractor period results in a consolidated waveform consisting of multiple peaks and troughs.

A

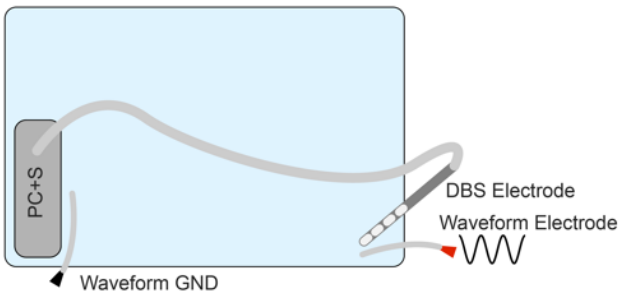


**Supplementary Figure 2: Experimental saline setup.** (a) The DBS lead and Activa PC+S case were immersed on opposite sides of a plastic container containing 1x phosphate buffered saline solution at room temperature. A platinum electrode connected to a waveform generator was placed adjacent to the stimulating electrode in order to simulate the neural signal appearing on the LFP. Single frequency oscillations were injected by the waveform generator alongside stimulation.

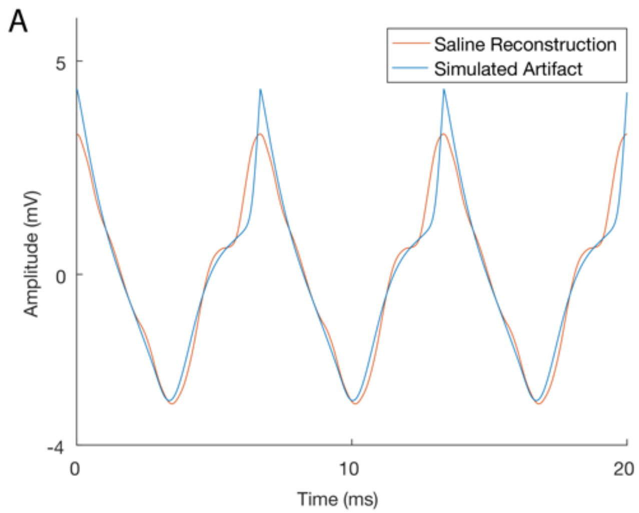


**Supplementary Figure 3: Simulated Activa PC+S output DBS waveform.** (a) Simulated DBS waveform pulse train (blue) and a PARRM reconstructed waveform pulse train from saline experiments (orange).

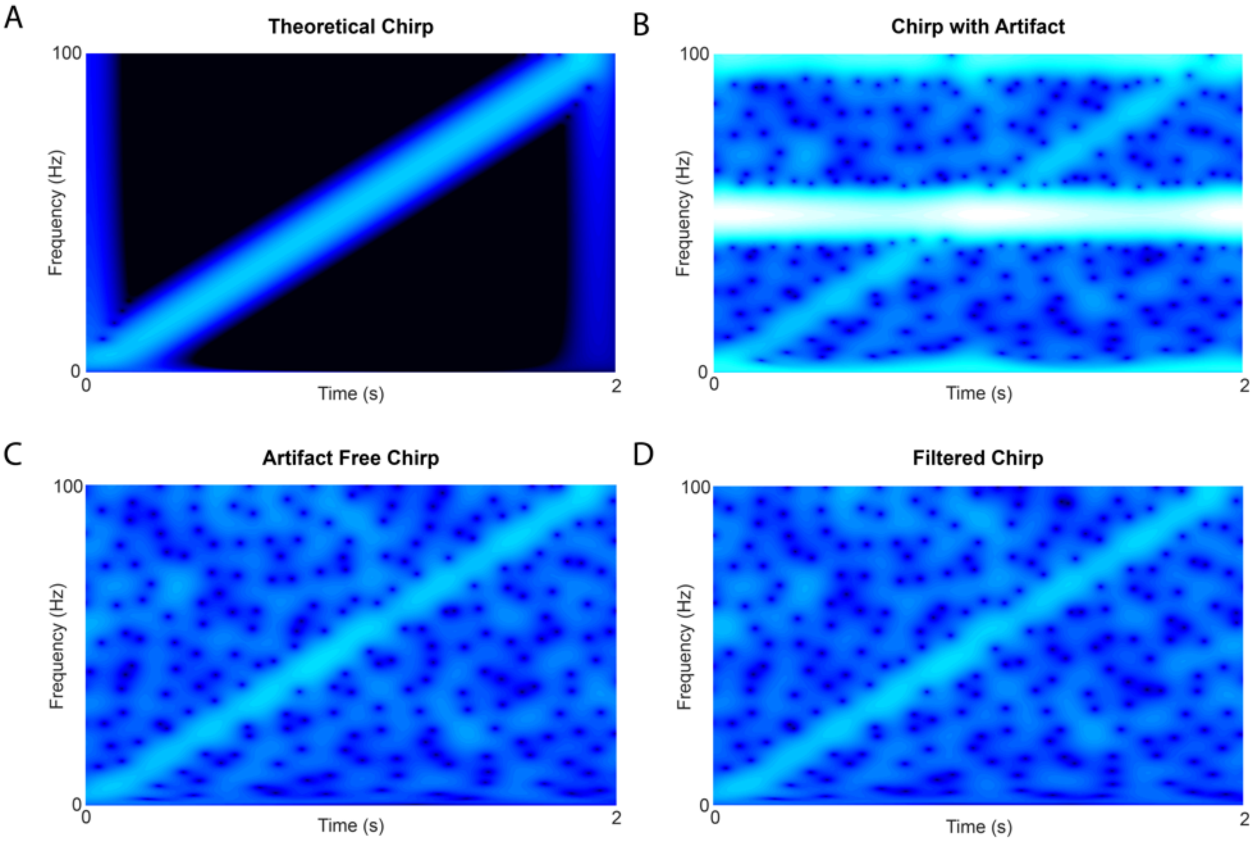


**Supplemental figure 4: Continuous wavelet transforms of simulated chirps.** (a) Continuous wavelet transform of a theoretical chirp with no noise or stimulation. (b) Continuous wavelet transform of a chirp with noise and stimulation at 150 Hz. (c) Continuous wavelet transform of a chirp with noise and no stimulation. (d) Continuous wavelet transform of a chirp with noise and stimulation at 150 Hz, filtered using PARRM.

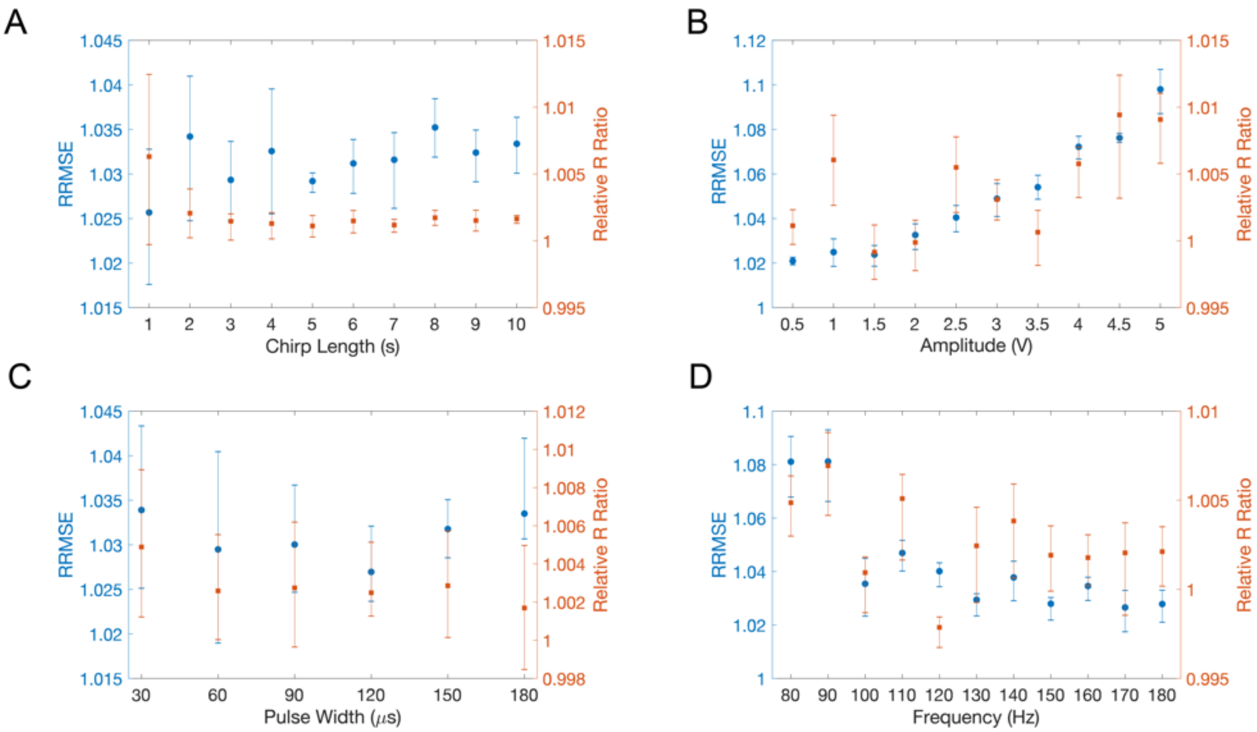


**Supplemental figure 5: Simulations show that PARRM is effective at a wide range of DBS parameters.** Relative RMSE (time domain) and R Ratio (frequency domain) for variable (a) chirp length, (b) amplitude, (c) pulse width, and (d) frequency. Error bars show 95% confidence interval on the median. Left Y axis in blue shows relative RRMSE. Right Y axis in orange shows Relative R Ratio.

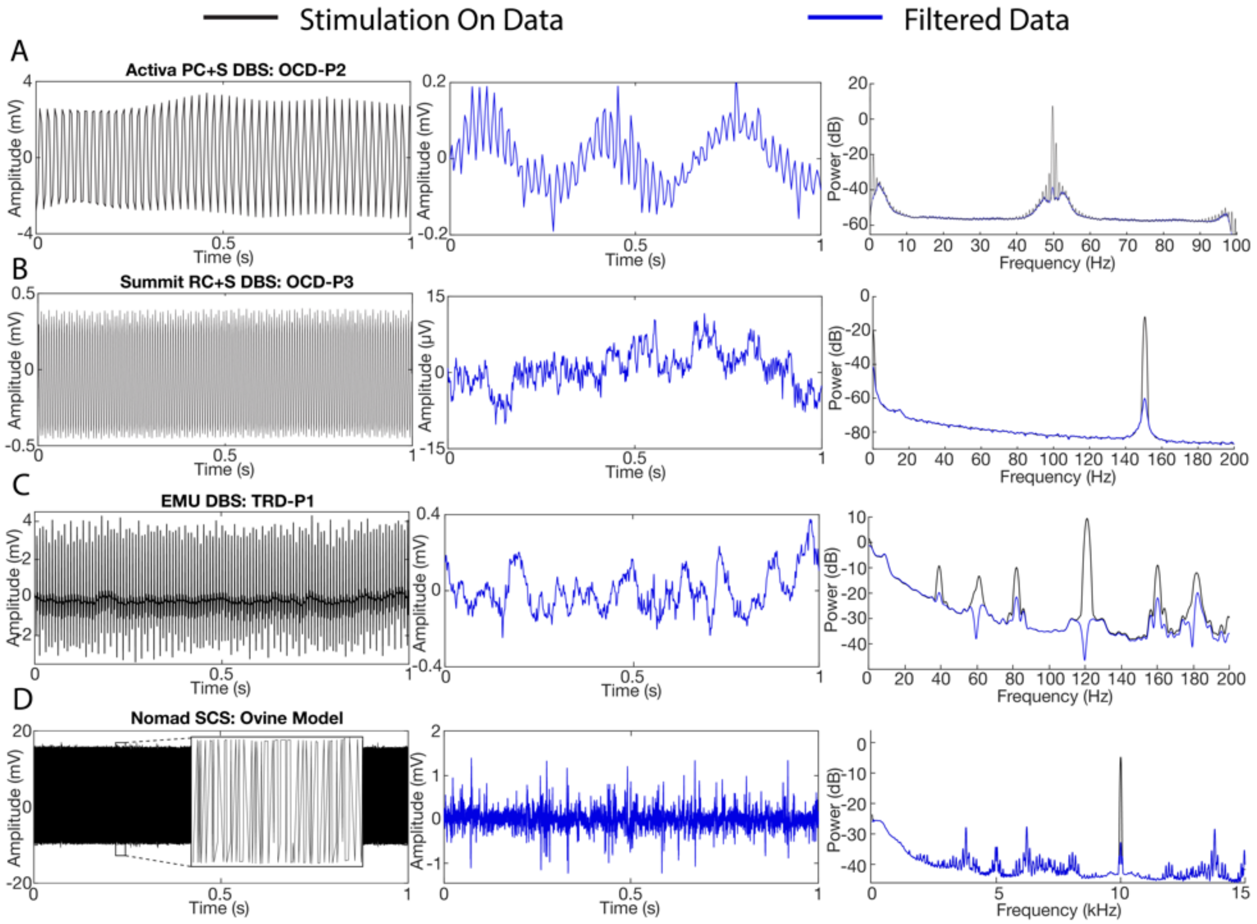


**Supplemental figure 6: Additional Demonstration of PARRM in human participants with DBS and SCS in ovine model.** (a-d) Raw time-voltage LFP trace, PARRM filtered time-voltage LFP trace, and average PSD before (black) and after (blue) PARRM filtering, collected during (a) 150 Hz stimulation sampled at 200 Hz using Activa PC+S in OCD-P2 left VC/VS, (b) 150.6 Hz stimulation sampled at 1000 Hz using Summit RC+S in OCD-P3 left VC/VS, (c) 120 Hz stimulation sampled at 2000 Hz in TRD-P1 right amygdala during a cognitive control task, (d) 10 kHz spinal stimulation sampled at 30 kHz in ovine model using Ripple Nomad

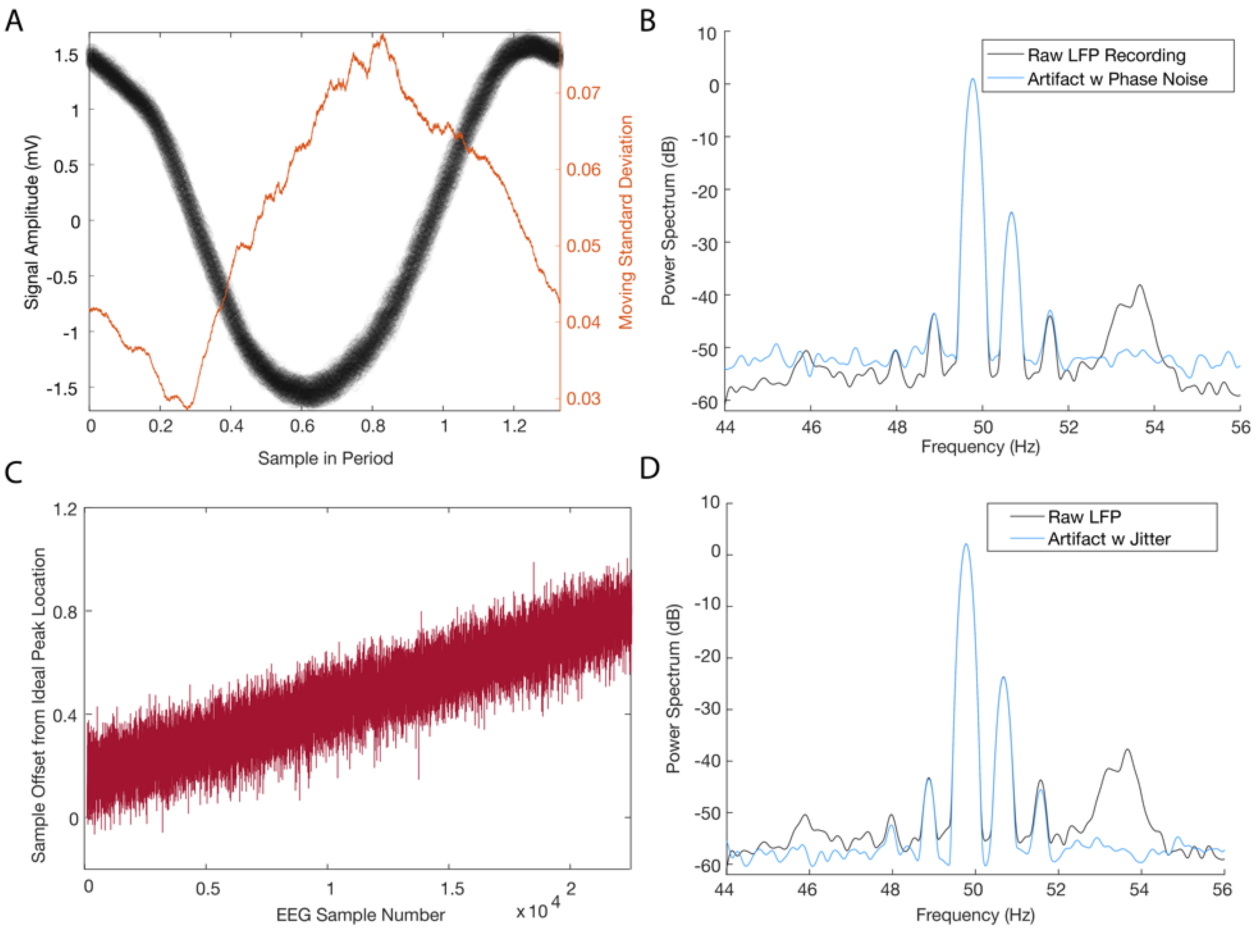


**Supplemental figure 7: Exploration of non-stationary oscillations leftover after PARRM in human data.** (a) Black points show LFP samples from a single human Activa PC+S recording overlaid on a single period. Orange line shows the 1000 sample moving standard deviation of the residuals after PARRM overlaid on a single period. (b) Black line shows the power spectral density of the raw LFP signal. Blue line shows the power spectral density of the reconstructed artifact with the addition of phase noise (gaussian noise with zero mean and standard deviation equal to the corresponding phase on the orange trace). (c) Trace showing deviation of true DBS pulse time in EEG from pulse time predicted by PARRM as it varies across a recording. (d) Black line shows the power spectral density of the raw LFP signal. Blue line shows the power spectral density of the reconstructed artifact with the addition of the jitter sequence from panel C for each pulse.

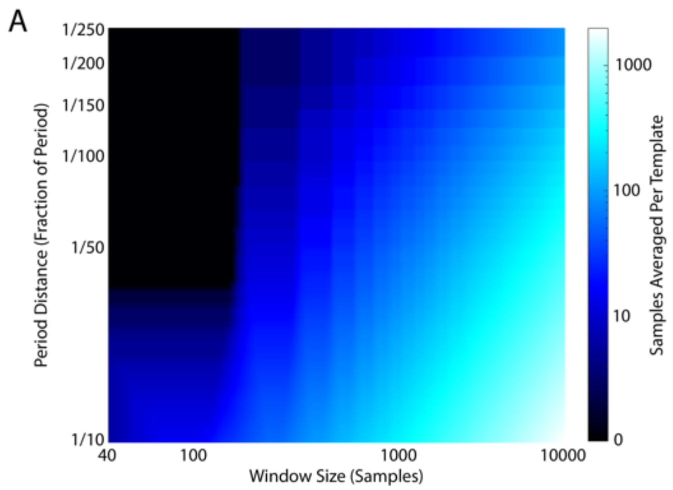


**Supplemental figure 8: Number of samples averaged as a function of window size and period distance.** (a) Heat map of the number of samples averaged as a function of period distance ($D_{period}$) a and half window size ($N_{bins}$). Darker blue indicates fewer samples averaged. Red point indicates the $D_{period}$ and $N_{bins}$ that were used for all analysis.

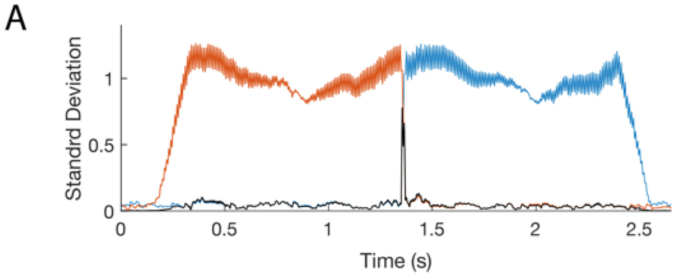


**Supplementary figure 9: Illustration of method for finding period jumps in LFP.** (a) Moving standard deviation (5 sample window) using prediction from the left (blue), right (orange), and product of moving standard deviation from the left and right (black) illustrate method for identifying period jump in LFP recording. The peak of the product signifies the location of the period jump in LFP, and is used for aligning LFP to the corresponding point in EEG.
